## Supplementary material for "Early Steps of Protein Disaggregation by Hsp70 Chaperone and Class B J-Domain Proteins are Shaped by Hsp110": Figure Supplements

### Supplementary Figures

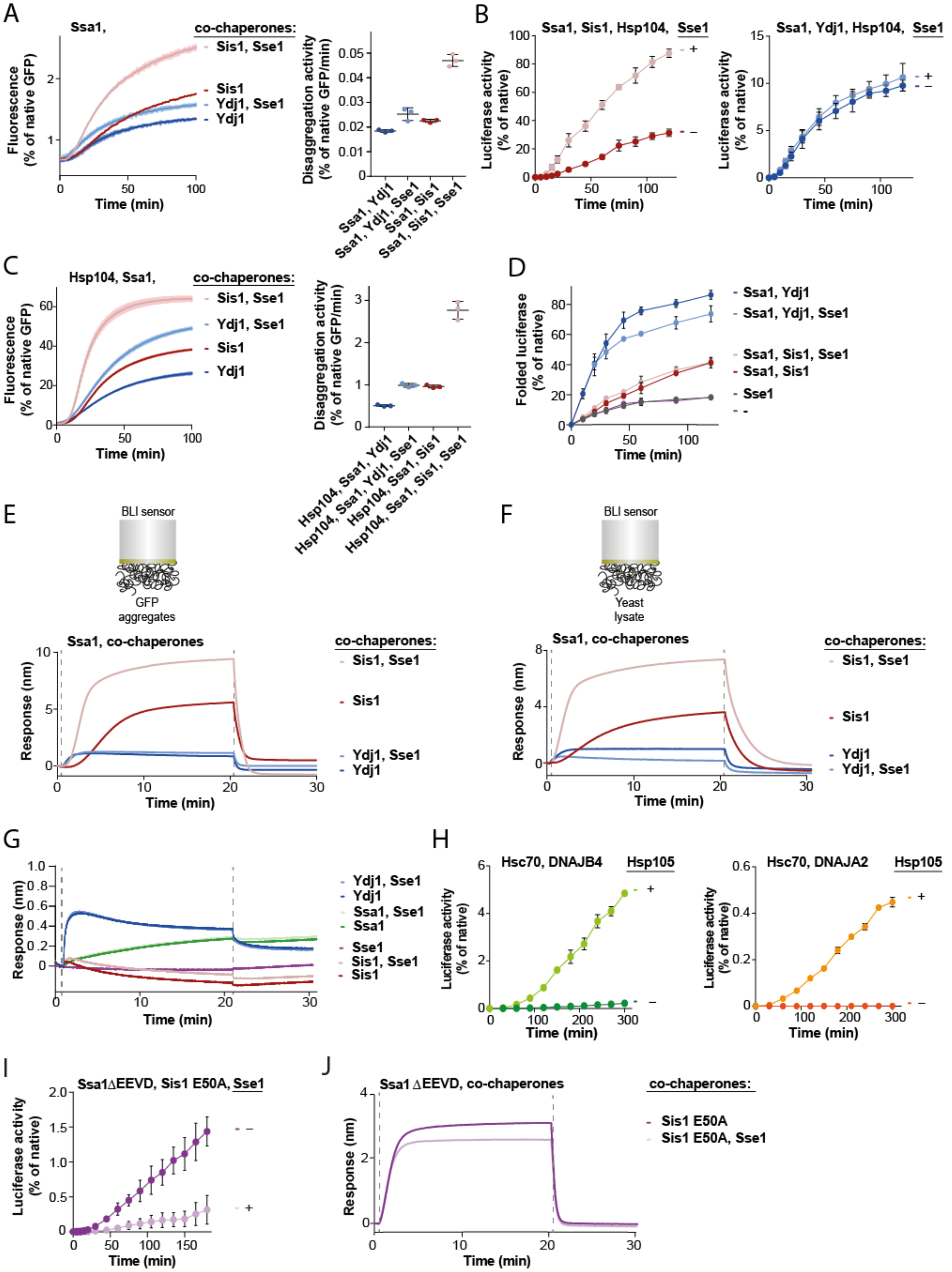

**Figure 1—figure supplement 1.** (A) Renaturation of heat-aggregated GFP by Ssa1-Sis1 +/- Sse1 or Ssa1-Ydj1 +/- Sse1 (1 μM Ssa1, 1 μM Sis1, 1 μM Ydj1 and 0,1 μM Sse1). (B) Refolding of aggregated luciferase by Ssa1-Sis1 +/- Sse1 (left) or Ssa1-Ydj1 +/- Sse1 (right) in the presence of Hsp104 (1 μM Ssa1, 1 μM Sis1, 1 μM Ydj1, 1 μM Hsp104 and 0.1 μM Sse1). (C) Renaturation of heat-aggregated GFP in the presence of Hsp104 with Ssa1-Sis1 +/- Sse1 or Ssa1-Ydj1 +/- Sse1 Hsp104 (1 μM Ssa1, 1 μM Sis1, 1 μM Ydj1, 1 μM Hsp104 and 0.1 μM Sse1). (D) Spontaneous folding of non-aggregated luciferase diluted from 5 M GuHCl, alone or assisted by the Hsp70 system (1 μM Ssa1, 1 μM Sis1, 1 μM Ydj1 and 0.1 μM Sse1). (E) Sensor covered with GFP aggregates or (F) yeast lysate incubated with Ssa1-Sis1 +/- Sse1 or Ssa1-Ydj1 +/- Sse1 (1 μM Ssa1, 1 μM Sis1, 1 μM Ydj1 and 0.1 μM Sse1). (G) Binding of the indicated combination of chaperones to luciferase aggregates immobilized on the BLI sensor (1 μM Ssa1, 1 μM Sis1, 1 μM Ydj1 and 0.1 μM Sse1). (H) Recovery of aggregated luciferase performed upon addition of human system comprising Hsc70-DNAJB4 +/- Hsp105 (left) or Hsc70-DNAJA2 +/- Hsp105 (right) (3 μM Hsc70, 1 μM DNAJB4, 1 μM DNAJA2 and 0.3 μM Hsp105). (I) Refolding of aggregated luciferase by Sis1 E50A and Ssa1 ΔEEVD +/- Sse1. (J) Binding of Sis1 E50A and Ssa1 ΔEEVD +/- Sse1 to luciferase aggregates immobilized on the BLI sensor (1 μM Ssa1 ΔEEVD, 1 μM Sis1 E50A and 0.1 μM Sse1). Shades and error bars represent SD from three independent repeats. The BLI experiments show a result representative for at least two independent repeats. Data from all the replicates are available in the Source Data File.

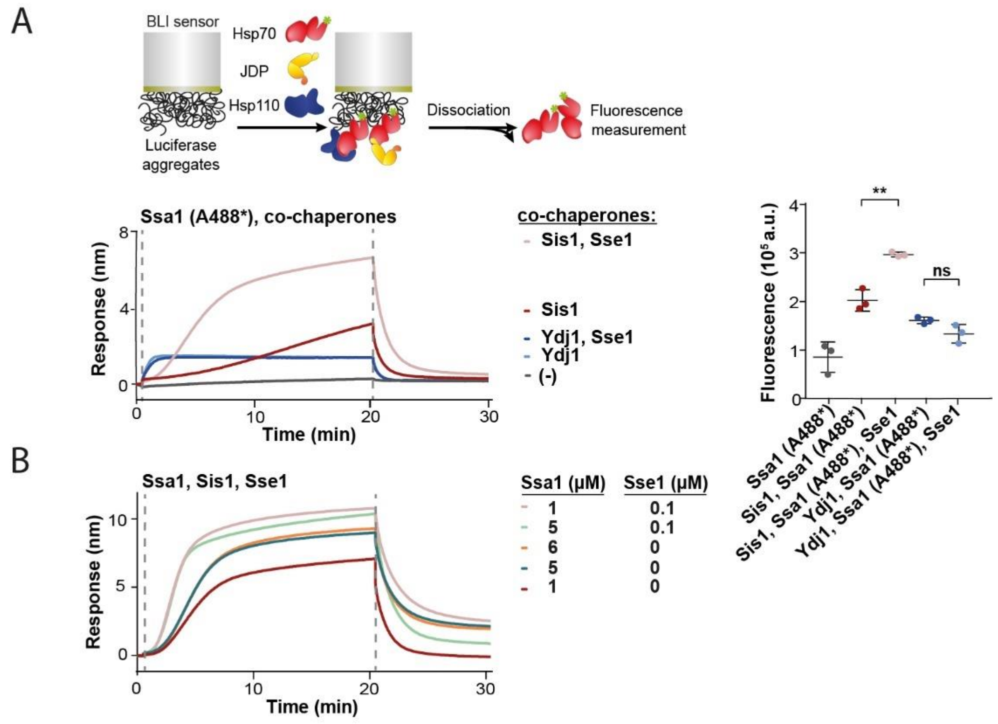


**Figure 1—figure supplement 2.** (A) Upper panel shows the scheme of the BLI experiment. Sensors covered with luciferase aggregates incubated with Ssa1 labelled with Alexa488 and Sis1 +/- Sse1 or Ydj1 +/- Sse1 (1 μM Ssa1 (A488*), 1 μM Sis1, 1 μM Ydj1 and 0.1 μM Sse1). Right panel shows fluorescence of Ssa1 (A488*) measured after the dissociation step. (B) Sensor-bound luciferase aggregates incubated with constant concentration of Sis1 (1 μM) and changing concentration of Ssa1 (1 μM, 5μM and 6 μM) +/- 0.1 μM Sse1.

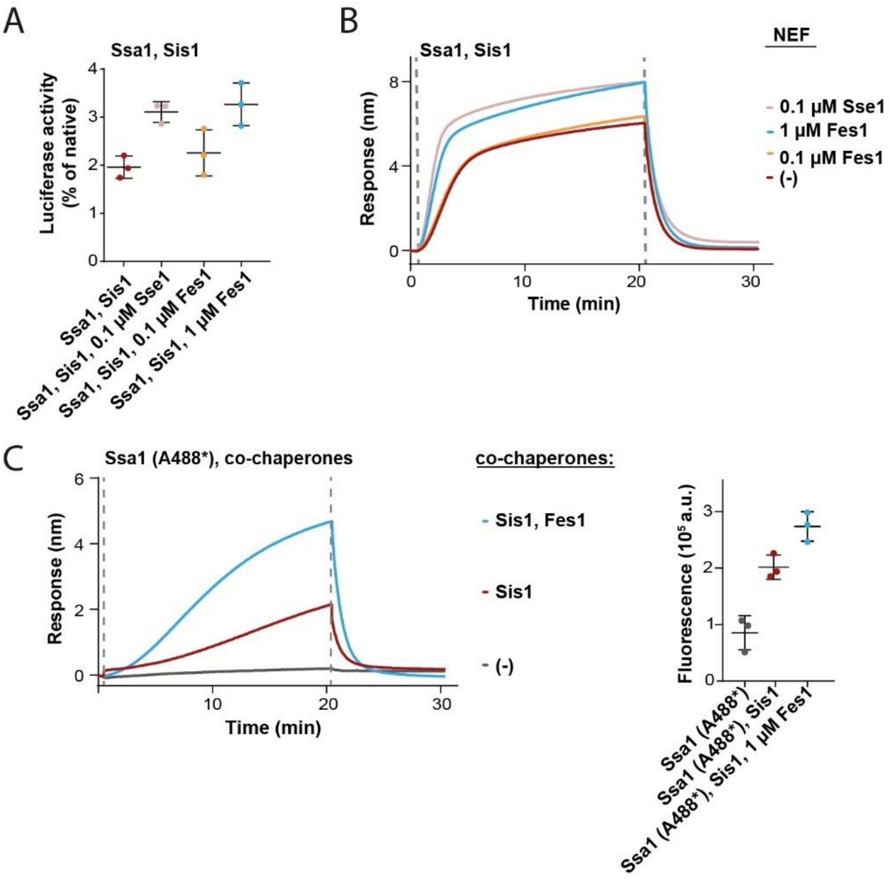


**Figure 1—figure supplement 3.** (A) Refolding of aggregated luciferase by Ssa1-Sis1 with 0.1 μM concentration of Sse1 or increasing concentrations of Fes1 measured after 1 h. (B) Sensor-bound luciferase aggregates incubated with Ssa1-Sis1 in the presence of the indicated concentrations Fes1 comparison to Ssa1-Sis1 with 0.1 μM Sse1. (C) Sensor covered with luciferase aggregates incubated with Ssa1 labelled with Alexa488 and Sis1 in the presence of 1 μM Fes1. Right panel shows fluorescence of Ssa1 (A488*) measured after the dissociation step. Binding kinetics together with fluorescence of Ssa1 (A488*) and Sis-Ssa1 (A488*) was adapted from Fig. S1G. Error bars show SD from three independent repeats. BLI curves show a result representative for at least two independent repeats. Data from all the replicates are available in the Source Data File.

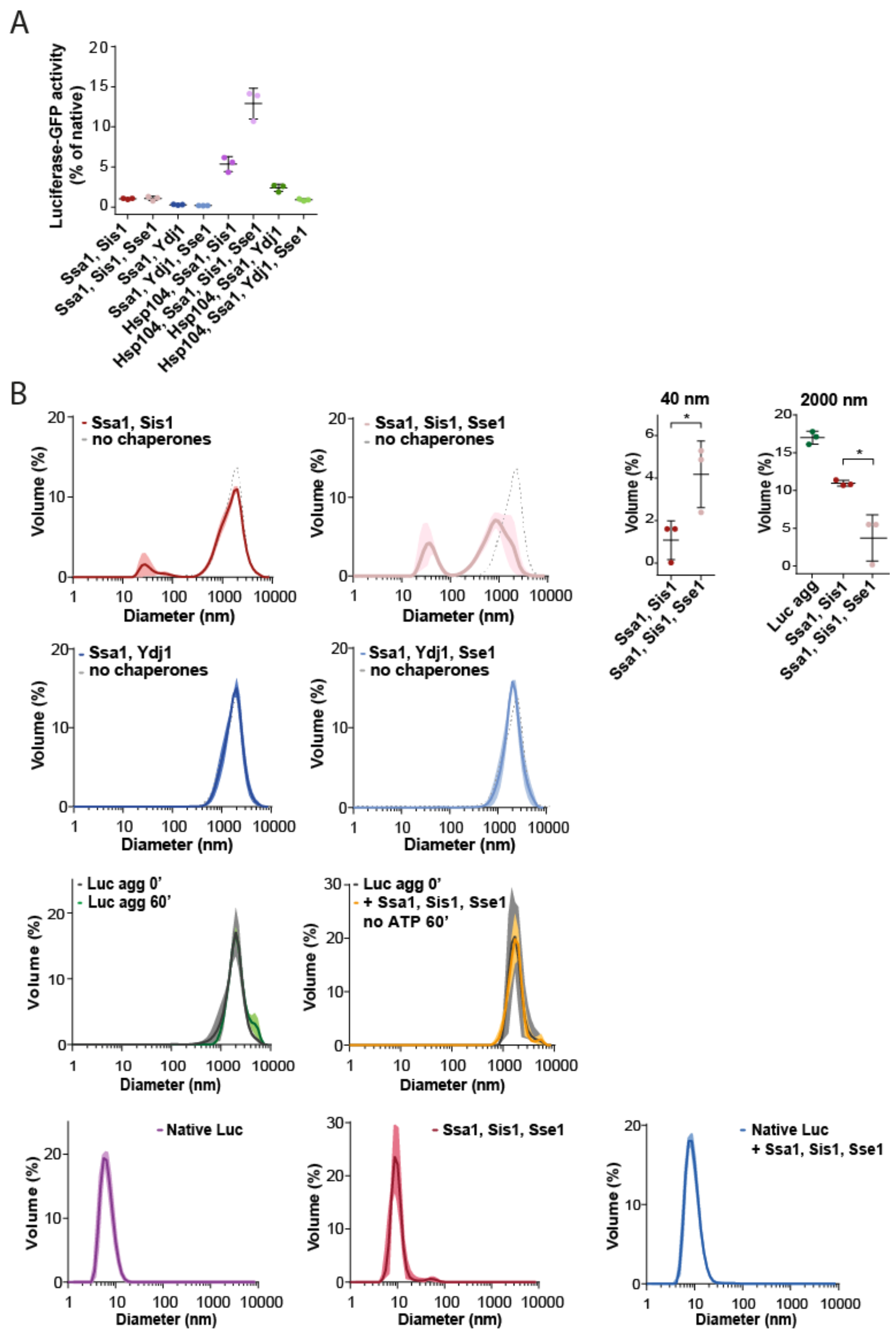


**Figure 2—figure supplement 1.** (A) Recovery of FLUC-EGFP aggregates (0.3 µM) by Hsp70 system with indicated combination of Ssa1 (1 µM), Sis1 (1 µM), Ydj1 (1 µM), Hsp104 (1 µM) or Sse1 (0.1 µM). L~~l~~uc-GFP activity was measured after 1 h incubation and normalized to the native activity. Error bars indicate SD from three repeats. (B) Size distribution measured with dynamic light scattering. Diameter of luciferase aggregates incubated alone or with Ssa1-Sis1 +/- Sse1 or Ssa1-Ydj1 +/- Sse1, as indicated in the figure, with ATP, unless stated otherwise, was measured after 1 h incubation. Proteins were used at the same concentrations as in A. Additionally, measurements with native luciferase, chaperones or chaperones with native luciferase were performed (lower panels). Lines are the average of three replicates

while the shades indicate standard deviation. Dashed lines designate the size distribution of the luciferase aggregates prior to the addition of chaperones. Upper right panel shows the height of the peak at 40 nm and 2000 nm for Ssa1-Sis1 with or without 0.1 μM Sse1, analyzed with the two-tailed t test: *p < 0,05, **p < 0,01 using the *GraphPrism* Software.

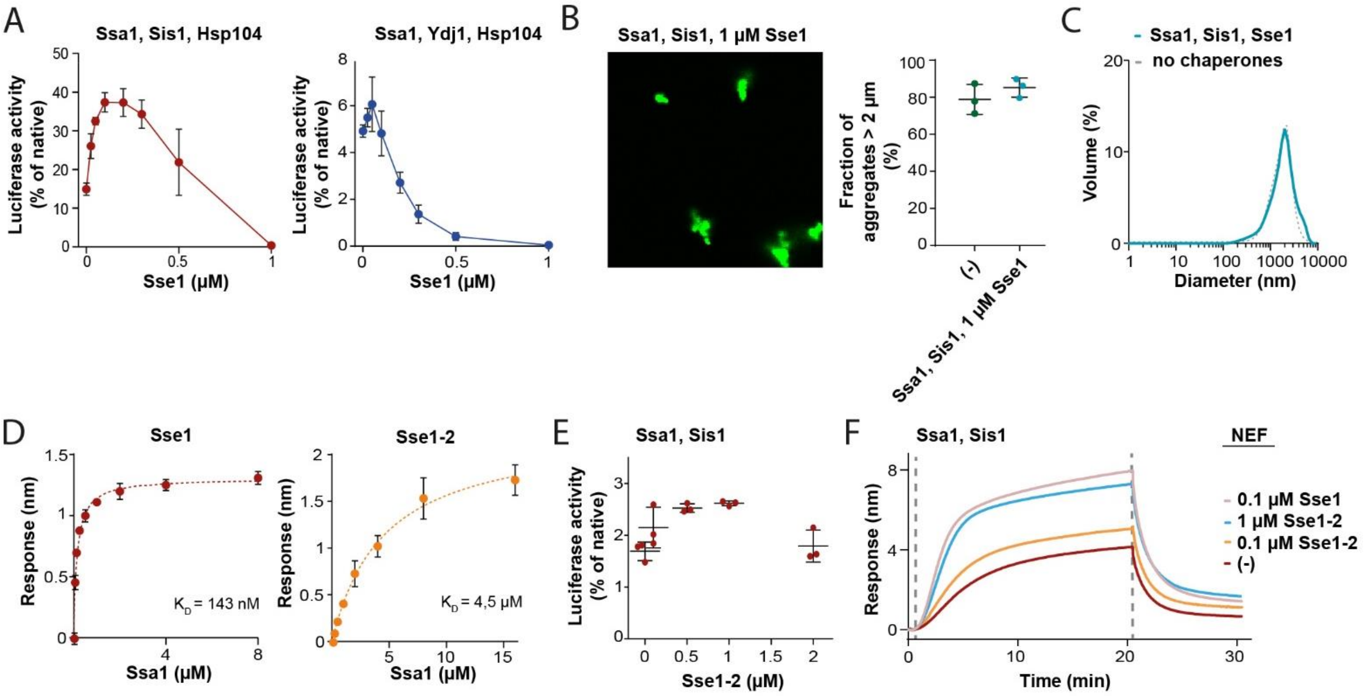


**Figure 3—figure supplement 1.** (A) Titration of Sse1 in the refolding of aggregated luciferase by Ssa1-Sis1 (red) or Ssa1-Ydj1 (blue) with Hsp104 (1 µM Ssa1, 1 µM Sis1, 1 µM Ydj1, 1 µM Hsp104 and indicated concentrations of Sse1). Activity of luciferase was measured after 1 h and normalized to the native activity. Shown is average and SD from three repeats. (B) Fluorescence microscopy images of luciferase-GFP aggregates incubated with Ssa1 (1 µM) and Sis1 (1 µM) in the presence of 1µM Sse1. Error bars show SD from three repeats. Quantification of the fraction of aggregates > 2 µm is from three independent replicates. Data for aggregates alone are from Figure 2B. (C) Size distribution of luciferase aggregates incubated with Ssa1 (1 µM) and Sis1 (1 µM) in the presence of 1 µM Sse1 measured by dynamic light scattering. (D) Dissociation constant is determined based on the level of Ssa1 binding to Sse1 or Sse1-2. Sse1 or Sse1-2 was immobilized on the BLI sensor through $His_6$-SUMO tag and Ssa1 was used at concentrations: 16 000, 8 000, 4 000, 2 000, 1 000, 500, 250, 125, 63 and 32 nM. Points indicate mean with SD from three independent experiments. The *One site – specific binding* model was fitted to the data with the *GraphPrism* software. (E) Titration of Sse1-2 in the refolding of luciferase aggregates assay with Ssa1-Sis1 (1 µM Ssa1, 1 µM Sis1). (F) Luciferase aggregates immobilized on the BLI sensor were incubated with Ssa1-Sis1 with either Sse1 or Sse1-2 at the indicated concentrations (1 µM Ssa1, 1 µM Sis1).

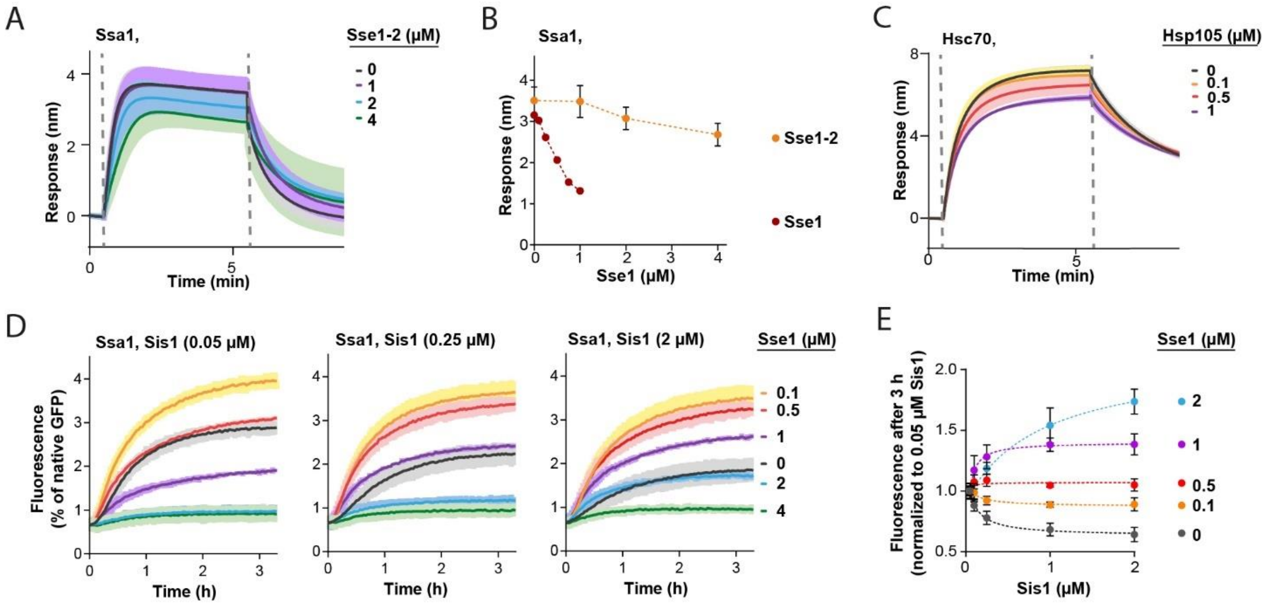


**Figure 4—figure supplement 1.** (A) Titration of Sse1-2 in the presence of 1 µM Ssa1 in the binding to the immobilized Sis1 on the BLI sensor. (B~~2~~) Comparison of the binding signal prior to the dissociation of Ssa1 in the presence of the indicated concentrations of Sse1 or Sse1-2. The binding signal of Ssa1-Sse1 WT was adapted from Fig. 4A. (C) Incubation of $His_6$-SUMO-DNAJB4 immobilized on the BLI sensor with Hsc70 and the indicated concentrations of Hsp105. (D) Titration of Sse1 in the GFP reactivation assay with Ssa1 (1 µM) and Sis1 at different concentrations: 0.05 µM, 0.25 µM, 2 µM. (E) Comparison of GFP fluorescence monitored after 3 h of incubation with Ssa1, Sis1 and Sse1. The values were adapted from Figure 4—figure supplement 1D and for 0.1 µM and 1 µM Sis1 they were adapted from Figure 4B. Dashed lines show fitting of the *[Agonist] versus response* model to the data from three experiments using the *GraphPrism* Software. Lines and points represent average values and error bars and shades indicate SD from three independent replicates.
